# Investigating the variability of physiological response functions across individuals and brain regions in functional magnetic resonance imaging

**DOI:** 10.1101/2024.06.13.596869

**Authors:** Laura B. Carlton, Georgios D. Mitsis, Michalis Kassinopoulos

## Abstract

Functional magnetic resonance imaging (fMRI) is a valuable neuroimaging tool for studying brain function and functional connectivity between brain regions. However, the blood oxygen level dependent (BOLD) signal used to generate the fMRI images can be influenced by various physiological factors, such as cardiac and respiratory activity. These physiological effects, in turn, influence the resulting functional connectivity patterns, making physiological noise correction a crucial step in the preprocessing of fMRI data. When concurrent physiological recordings are available, researchers often generate nuisance regressors to account for the effect of heart rate and respiratory variations by convolving physiological response functions (PRF) with the corresponding physiological signals. However, it has been suggested that the PRF characteristics may vary across subjects and different regions of the brain, as well as across scans of the same subject. To investigate the dependence of PRFs on these factors, we examine the performance of several different PRF models, in terms of BOLD variance explained, using resting-state fMRI data from the Human Connectome Project (N=100). We examined both one-input (heart rate or respiration) and two-input (heart rate and respiration) PRF models and show that allowing PRF curves to vary across subjects and brain regions generally improves PRF model performance. For one-input models, the improvement in model performance gained by allowing spatial variability was most prominent for respiration, particularly for a subset of the subjects (about a third) examined. Allowing for subject-specific or regional variability in the cardiac response function resulted in a significant model performance improvement only when using a two-input PRF model. Overall, our results highlight the importance of considering spatial and subject-specific variability in PRFs when analyzing fMRI data, particularly regarding respiratory-related fluctuations.

## 1. Introduction

Functional magnetic resonance imaging (fMRI) has emerged as a valuable tool for studying brain function and understanding neurological diseases. This imaging technique relies on metabolic changes triggered by changes in neural activity within the brain. When a brain area is activated, it requires more energy for neural signaling, increasing local cerebral metabolism. In turn, this triggers a series of events for the delivery of additional energy to the activated brain regions. Specifically, the amount of oxygen being delivered to these regions is increased via an increase in cerebral blood flow (CBF), which also increases the concentration of oxyhemoglobin in the blood. Oxyhemoglobin and deoxyhemoglobin have different magnetic properties and their relative concentrations determine the contrast in fMRI images (Glover, 2011; Ogawa et al., 1990). Due to this, the blood oxygen level dependent (BOLD) contrast mechanism is typically used in fMRI as an indirect measure of neural activity.

Many researchers have focused on using resting-state fMRI scans to identify and study resting-state networks (RSNs) in the brain. These functional networks include regions of the brain that exhibit similar low frequency oscillations (<0.15Hz) (Biswal et al., 1995; Fox & Raichle, 2007; Smith et al., 2009). RSNs can provide insight into the functional connectivity of the brain but they need to be interpreted carefully, as the BOLD signal can be affected by physiological variables such as heart rate and respiration, which may cause spurious results in the analysis of functional connectivity (Birn et al., 2006; Murphy et al., 2013; Shmueli et al., 2007; Tong et al., 2019; Xifra-Porxas et al., 2021). Heart rate fluctuates due to the action of the autonomic nervous system causing changes in CBF are, in turn, reflected on the fMRI signal (Murphy et al., 2013; Tong et al., 2019). The movement of the chest during respiration can cause changes in the magnetic field within the scanner which can shift the brain image (Raj et al., 2001). Furthermore, changes in breathing rate (BR) and depth can affect the concentration of arterial CO2, a potent vasodilator and, thus, affect CBF (Birn et al., 2006; Wise et al., 2004). It has also been shown that spontaneous arterial CO2 variations cause low frequency fluctuations in the BOLD signal that can be misinterpreted as neural activity (Prokopiou et al., 2019; Stickland et al., 2021; Wise et al., 2004). Finally, arterial blood pressure also fluctuates over time and these variations influence fMRI timeseries through their effects on CBF (Whittaker et al., 2019). Thus, being able to remove these physiological artefacts is important for accurately quantifying the fMRI signatures of the underlying neural activity.

Many models have been proposed to effectively remove physiological noise; however, a conclusion has yet to be made on the best practice to achieve this. A balance must be reached whereby most of the physiological artefacts are adequately removed without removing a significant fraction of the neural signal of interest. An additional challenge is that there is often a temporal coupling of physiological fluctuations and neural activity that is hard to disentangle.

To mitigate the effects of physiological confounds, peripheral recordings obtained simultaneously with fMRI scans have been shown to be very useful. Birn et al. (2008) and Chang et al. (2009) first proposed the use of linear convolution models to create nuisance regressors, which can be incorporated in the general linear model (GLM) framework typically used in fMRI studies to correct for physiological confounds. According to these models, the trace of heart rate (HR) is convolved with a cardiac response function (CRF) to generate cardiac-related nuisance regressors. Similarly, respiration volume per time (RV), which is a measure of respiration depth and rate, is convolved with the respiratory response function (RRF) to create a regressor that accounts for changes in the BOLD signal caused by variations in respiratory patterns. These physiological regressors can then be removed from the BOLD signal through linear regression. These models have been widely used for preprocessing of fMRI data, but they are limited in their ability to account for variability in physiological responses across subjects and brain regions.

Recent work has examined whether using only one invariant physiological response function (PRF) curve for noise removal is appropriate due to the spatial and subject variability in the brain’s response to physiological fluctuations (Chen et al., 2020; Falahpour et al., 2013; Kassinopoulos & Mitsis, 2019). The brain vasculature is heterogeneous across brain regions, as well as across subjects (Bernier et al., 2018), and this is likely to affect the dynamics of region-specific responses to changes in HR and breathing rate (BR). Kassinopoulos and Mitsis (2019) reported that the best model performance, in terms of BOLD signal variance explained, was achieved when subject variability was accounted for in the PRF curves, whereas no additional improvement was observed when allowing for regional variability. However, the findings by Chen et al. (2020) provided evidence in favor of accounting for spatial variability in the RRF curves. This discrepancy may be related to the fact that these two studies employed models with a different number of basis functions, and thus flexibility. Given this, the aim of the present study was to rigorously assess the benefit of accounting for subject and regional variability in the context of PRF estimation, using both one-input (heart rate or respiration) and two-input (heart rate and respiration) PRF models. To this end, we tested seven linear convolution models with varying degrees of flexibility at three different spatial resolutions (voxel, parcel and whole-brain level). Our findings suggest that allowing subject-specific variability for the RRF curves is beneficial for noise removal at all levels of spatial resolution. Moreover, in approximately a third of subjects, regionally variable RRF curves were found to explain additional variance in both voxel- and parcel-wise analysis. When considering only cardiac models, we did not observe a significant improvement when allowing for subject or regional variability in the CRF curves. However, when considering two-input models, allowing for subject and regional variability was found to yield the best performance.

## 2. Methodology

### 2.1. Human Connectome Project (HCP) Dataset

The resting-state fMRI data used in this study are from the S1200 release of the 3T HCP dataset which consists of young, healthy individuals (age range: 22-35 years) (Glasser et al., 2016; Smith et al., 2013; Van Essen et al., 2013; Van Essen et al., 2012). The data was acquired on two different days. On each day two 15 min scans were collected with the left-to-right (LR) and right-to-left (RL) phase encoding direction. During each fMRI scan, 1200 frames were acquired using a gradient-echo echo-planar imaging (EPI) sequence with a multiband factor of 8, spatial resolution of 2 mm isotropic voxels and a TR of 0.72 s (Glasser et al., 2016). Further details of the data acquisition parameters can be found in previous publications (Smith et al., 2013; Van Essen et al., 2012).

Cardiac and respiratory signals were measured simultaneously with the fMRI scans. This was done using a standard Siemens pulse oximeter placed on the fingertip and a breathing belt placed around the chest with a 400 Hz sampling rate. We only considered subjects with available data from all four scans, and excluded subjects based on the quality of the physiological recordings. Pulse oximeter and breathing belt signals from ∼1000 subjects were first visually inspected to determine their quality, since their traces are often not of sufficient quality for reliable peak detection (Power, 2019). The selection process resulted in a final dataset from 392 subjects, of which 100 were used in the present study (ID numbers provided in Supp. Material). Briefly, the minimal initial preprocessing pipeline included removal of spatial distortion, motion correction via volume realignment, registration to the structural image, bias-field correction, 4D image intensity normalization by a global mean, brain masking, and non-linear registration to MNI space.

### 2.2. Preprocessing

The HR and preprocessed respiratory signals extracted in Xifra-Porxas et al. (2021) were used in the present study. The pulse wave from the oximeter was processed to automatically detect beat-to-beat intervals (RR). The HR signal was computed as the inverse of the time difference between pairs of adjacent peaks and converted to units of beats-per-minute (bpm). HR signals were visually checked to ensure that no abnormalities and outliers were present. If spurious changes in HR were present due to sporadic noisy cardiac measurements, an outlier replacement filter was used to eliminate them.

The signal from the breathing belt was linearly detrended, visually inspected and corrected for outliers using a replacement filter. It was subsequently low-pass filtered at 5 Hz and z-scored to set the mean to zero and the variance to one. Respiratory variation (RV), defined as the standard deviation of the movement of the respiration bellows in a 6 sec sliding window, was extracted from the breathing belt signal. Both the HR and RV data were temporally filtered with a high-pass filter at a cut off frequency of 0.008 Hz and z-scored. The signals were then re-sampled at 10 Hz.

The raw fMRI volumes were spatially smoothed with a Gaussian filter of 5 mm full width half maximum (FWHM). The data were linearly detrended, temporally filtered with a high-pass filter at a cut off frequency of 0.008 Hz and normalized. Nuisance regressors were then removed through linear regression. The regressors included the six demeaned and detrended motion parameters and their derivatives, as well as 6^th^ order RETROICOR regressors for cardiac artefacts and the 3^rd^ order RETROICOR regressors for respiratory artefacts using the physiological recordings at a sampling rate of 400 Hz (Glover et al., 2000).

For the analysis at the parcel level, the Gordon Atlas was used to parcellate the brain into 333 distinct regions (Gordon et al., 2016). The BOLD signals from all voxels within a parcel were averaged to get parcel-specific BOLD timeseries. The global signal (GS) was calculated by taking the mean signal from all the voxels. All the data were high-pass filtered at a cut off frequency of 0.008 Hz and z-scored. The first 40 volumes of each timeseries at the voxel, parcel and whole-brain level were removed, while the corresponding physiological signals were retained to account for the duration of the effects of HR and respiratory variations.

### 2.3. Physiological response functions

This study examined seven linear convolution models outlined in Table 1. The standard PRF curves and the employed basis functions are shown in Figure 1. Each model contained a cardiac response function (CRF) convolved with HR to model the cardiac related effects on the BOLD signal and a respiratory response function (RRF) convolved with RV to model the respiratory related effects on the BOLD signal. These convolutions generated physiological regressors associated with cardiac- and respiratory-related fluctuations, i.e.:

**Table 1.**
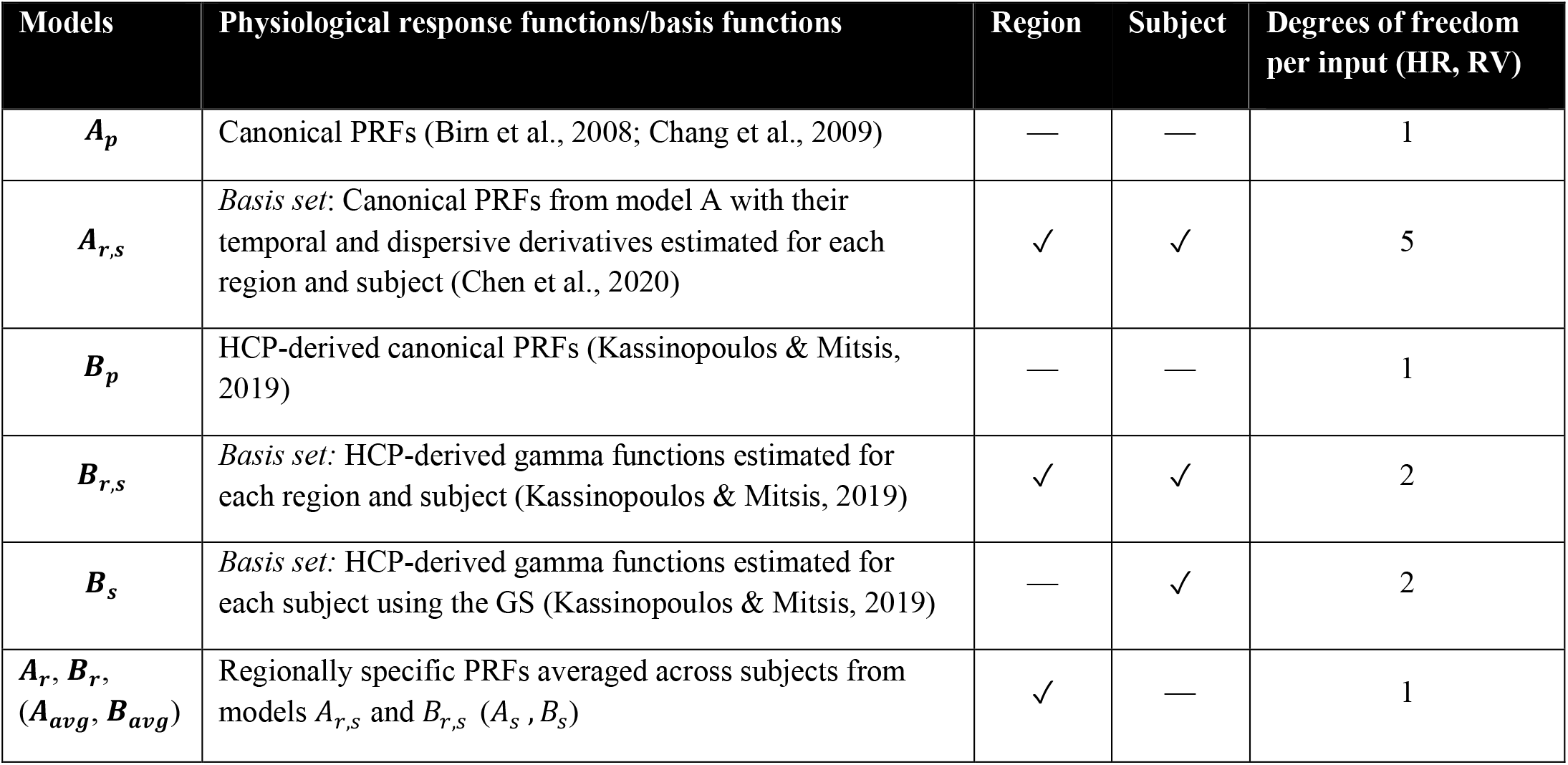
Overview of the 7 PRF models. The subscript *p* denotes a PRF estimated from a population in a previous study (Birn et al., 2008; Chang et al., 2009; Kassinopoulos & Mitsis, 2019), *r* denotes a regionally variable model, and *s* denotes a subject-specific model. Models denoted by A are based on the PRFs of Birn et al. (2008) and Chang et al. (2009), while models denoted by B are based on the PRFs of Kassinopoulos and Mitsis (2019).

**Figure 1.**
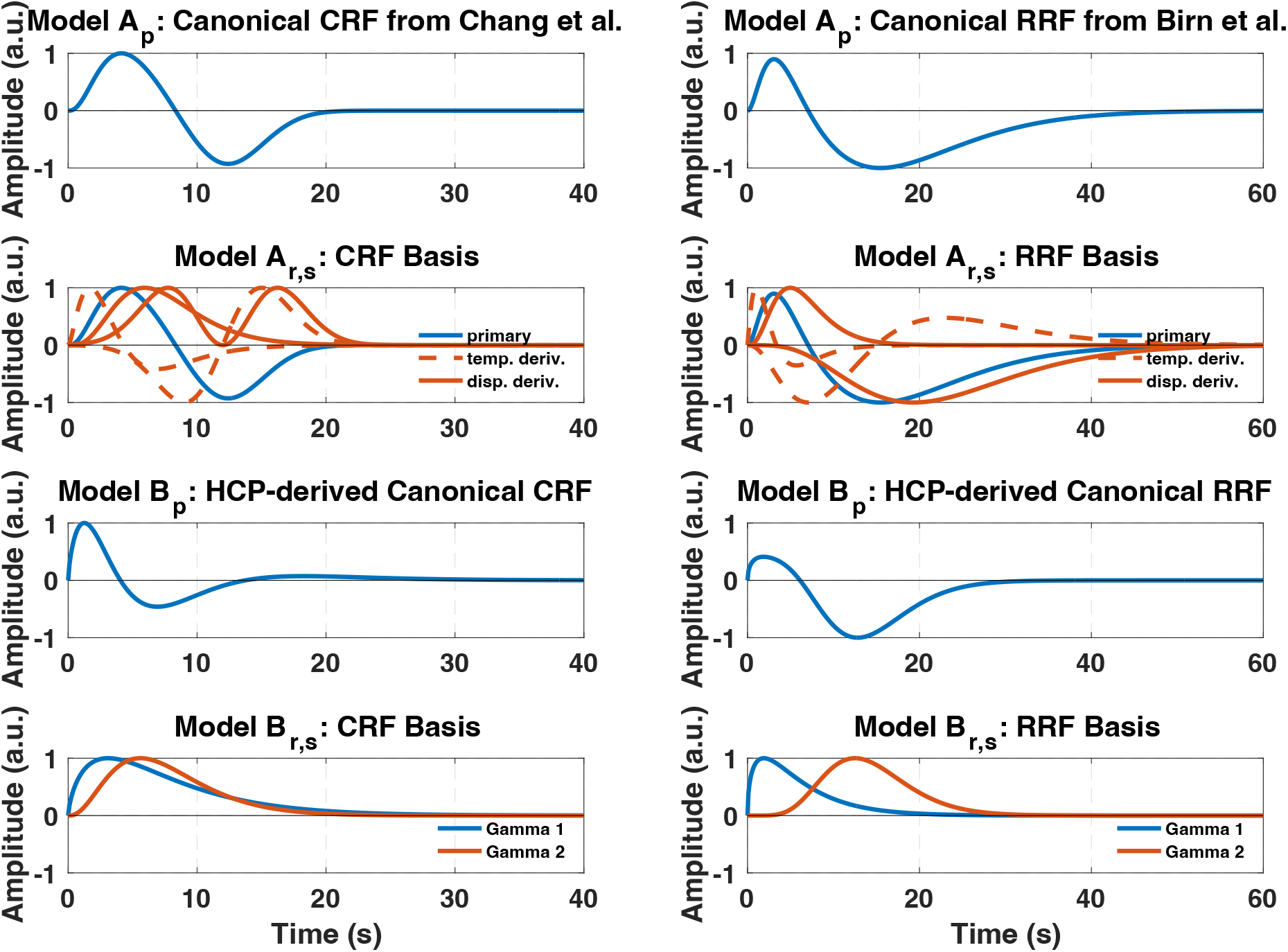
(first row) Model *A_p_* employs one invariant curve for the CRF and RRF based on the work of Chang et al. (2009) and Birn et al. (2008), respectively. (second row) The basis set of model *A_r,s_* is from the work of Chen et al. (2020). The canonical curves are shown in blue. The temporal derivatives are indicated with a dashed orange line and the dispersive derivatives are shown with a solid orange line. (third row) The canonical curves for model *B_p_* are based on the work of Kassinopoulos and Mitsis (2019). (fourth row) The basis for model *B_r,s_* is derived by taking the individual gamma functions from the PRFs of model *B_p_*.

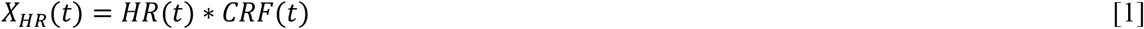

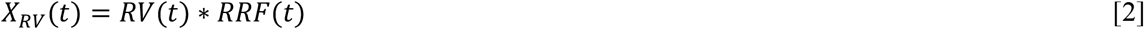

The regressors *X_HR_* and *X_RV_* were then downsampled to the fMRI sampling rate and high-pass filtered at a cut off frequency of 0.008 Hz. The examined models had varying degrees of flexibility and were applied to the voxel, parcel and whole-brain (GS) fMRI timeseries.

Certain models required the estimation of the PRFs on a regional and subject level. In these cases, the basis expansion technique was used to estimate the PRF curves using a suitable basis set 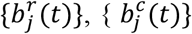 where the final PRF curve was obtained as a weighted sum of the selected *L* basis functions (superscripts *r* and *c* correspond to cardiac and respiratory basis sets and weights respectively):

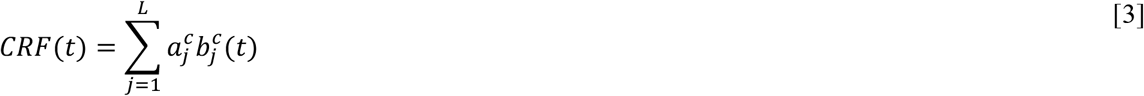

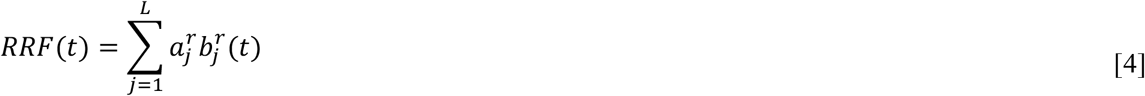

The expansion coefficients, 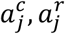, were determined through ordinary least squares fitting-based regression between the HR and RV signals convolved with the basis functions and the BOLD signal as follows:

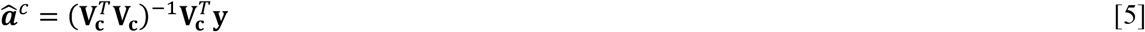

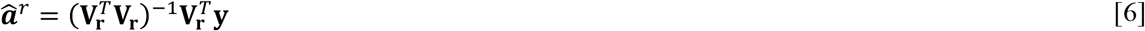

where **y** corresponds to the BOLD time series and the regressor matrices **V_c_**, **V_r_** contain the convolution values between the HR and RV signals with the basis functions *b_j_*(*t*) at different time points.

#### 2.3.1. Model *A_p_* – Standard canonical PRFs

Model *A_p_* employed the invariant CRF and RRF curves proposed by Chang et al. (2009) and Birn et al. (2008), respectively. The RRF was a weighted sum of two gamma functions while the CRF was a weighted sum of a gamma and a Gaussian function. The PRFs are expressed as follows:

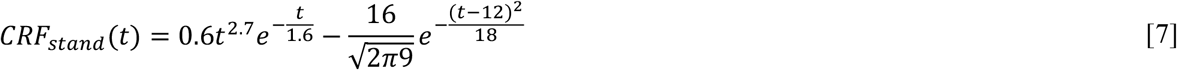

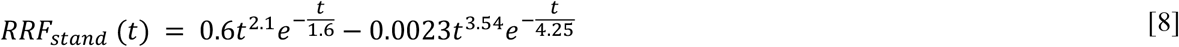

#### 2.3.2. Model *A_r,s_* – Standard canonical PRFs with their temporal and dispersive derivatives

Based on the work of Chen et al. (2020), model *A_r,s_* used the standard PRFs from model *A_p_*, as well as their temporal and dispersive derivatives to generate a basis set of five functions for both the CRF and RRF. The basis functions are given by the following relations:

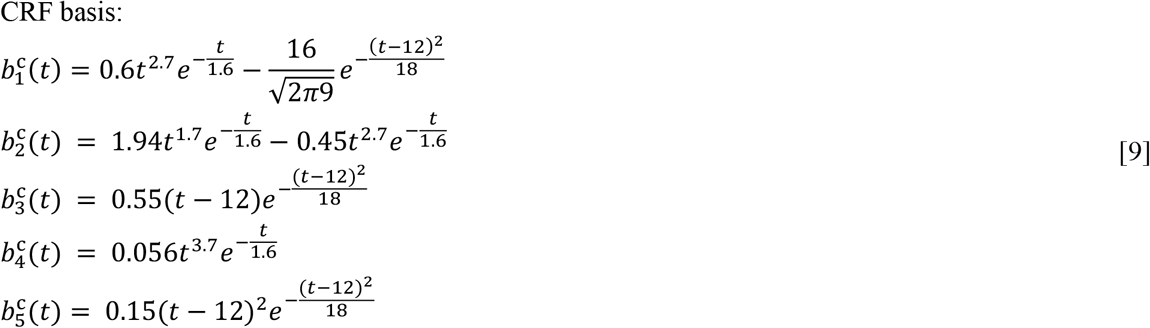

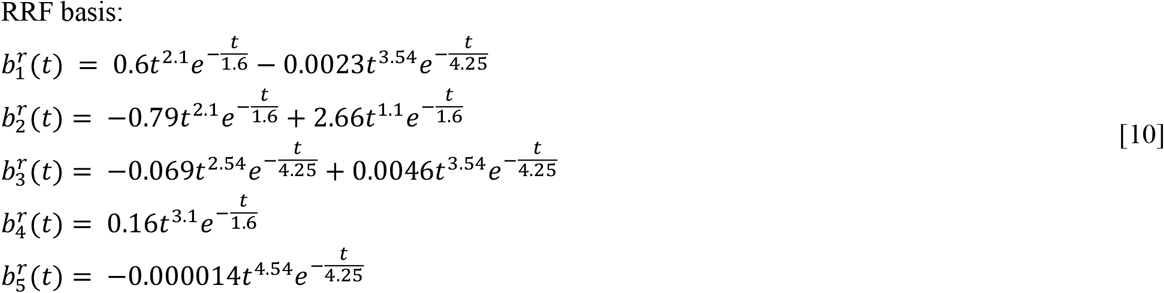

Each basis function was convolved with the appropriate physiological signal to generate a set of five cardiac and five respiratory regressors. These regressors were then fitted to the BOLD signal using linear regression as described above (Eq 5,6). The PRF curves were generated by taking the weighted sum of the five basis functions using the expansion coefficients estimated through the linear regression as the weighting coefficients (Eq 3,4).

#### 2.3.3. Model *B_p_* – HCP-derived PRFs

Model *B_p_* used the population-specific PRFs, *CRF_pop_* and *RRF_pop_*, reported in Kassinopoulos and Mitsis (2019). These were estimated from a subset of subjects (N=41) from the HCP dataset using a novel algorithm based on basis expansions with double-gamma functions. The HCP-derived PRFs can be expressed as follows:

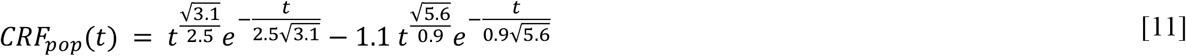

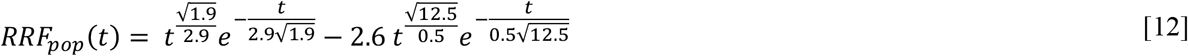

#### 2.3.4. Model *B_r,s_* – HCP-derived gamma functions

To allow subject and spatial variability, model *B_r,s_* used the individual gamma functions of model *B_p_* without their weighting coefficients as basis functions, expressed by the following relations:

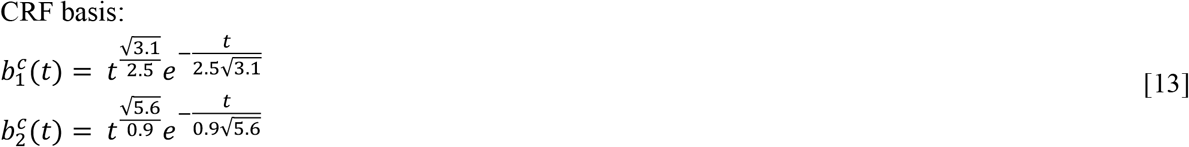

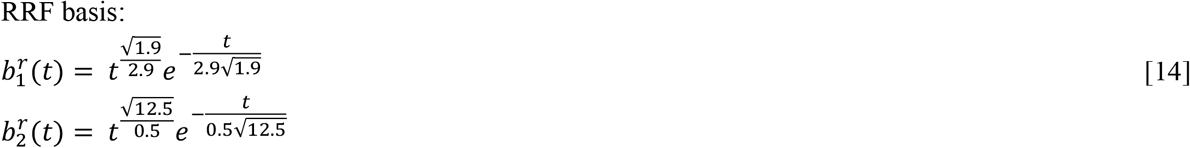

Each basis function was convolved with the corresponding physiological signal to generate either cardiac or respiratory regressors. These regressors were then fitted to the BOLD signal using linear regression as described above (Eq 5,6). The PRF curves were generated by taking the weighted sum of the five basis functions using the expansion coefficients estimated through the linear regression as the weighting coefficients (Eq 3,4).

#### 2.3.5. Model *A*_*s*_/*B*_*s*_ – subject specific PRFs

To obtain PRFs that were subject-but not region-specific, the GS was used to estimate them using the basis functions considered in models *A_r,s_* and *B_r,s_*. Specifically, a single CRF and RRF curve was generated for each subject by using the expansion coefficients estimated through linear regression (Eq 5,6) when fitting the HR and RV signals convolved with the basis functions to the GS.

#### 2.3.6. Model *A*_*r*_/*B*_*r*_ (*A_avg_*/*B_avg_*)

The regional-specific PRF curves in models *A_r_* and *B_r_* were generated by averaging the regionally specific curves obtained with *A_r,s_* and *B_r,s_* across all subjects. Similarly, at the whole-brain level, the models *A_avg_* and *B_avg_* were generated by averaging the curves from models *A_s_* and *B_s_* across all subjects.

### 2.4 Model evaluation

Model performance was evaluated using the Pearson correlation coefficient between the BOLD timeseries and the model output. Cross validation was implemented for the models that required PRF estimation (i.e. *A_r,s_, B_r,s_, A_s_, B_s_*). Specifically, the PRFs were estimated with the basis expansion technique described above using the first scan of a session from each subject and the model performance was evaluated on the second scan of that session. For models that did not require PRF estimation (i.e. *A_p_, B_p_, A_r_, B_r_*), the performance was evaluated directly on the second scan. The correlation was determined for the cardiac and respiratory regressors for each input (HR and RV) separately and for both inputs combined. A pairwise t-test was used to compare the performance across the different models.

At the voxel level, the evaluation of model performance was restricted to voxels for which the correlation values for the two-input model were in the top 5%, so that only regions strongly affected by variations in heart rate and respiratory patterns were considered for model comparison. At the parcel level, all parcels were considered for model comparison, as the averaging of voxel timeseries within a parcel tends to suppress random noise and enhance the effects of physiological processes.

### 2.5. Assessment of regional and subject variability

Based on the results of model comparisons (Section 3.1), further analysis was done to identify the subjects for which using region-specific RRFs significantly improved performance. All four scans from 100 subjects from the HCP dataset were used. First, the improvement of model *B_r_* as compared to model *B_p_* was assessed by computing the parcel-wise differences in the correlation values between the respiratory-related model output and BOLD signal obtained by the two models. K-means clustering was subsequently performed to segregate the scans into two groups using the parcel-wise differences. This step categorized the scans into a group that exhibited improvement when regional variability was allowed and a group that did not (designated the ‘improvement’ group and the ‘no improvement’ group).

To quantify whether certain subjects benefitted more from allowing PRF regional variability, a subject specificity score was used. Specifically, each subject was assigned a value based on the number of scans that were categorized in the same group. Subjects who had all scans in the same group were assigned a value of 1, subjects with one scan in one group and three scans in the other were assigned a value of 0.5, and subjects with two scans in one group and two in another were assigned a value of 0. The value assignment was the same regardless of which groups the scans belonged to. This metric indicates the degree to which the subjects had all their scans in one group. For example, for a subject with all four scans in the same group the overall score would be 1, indicating a strong degree of subject specificity in the clustering. Conversely, for subjects with two scans in each group the score would be zero indicating a lack of subject specificity in the clustering.

To compute an overall score, the subject-specific score values were averaged across all subjects. Subsequently, the significance of the subject specificity score was assessed by comparing its distribution to a null distribution generated by permutating the data and computing the subject specificity score 10,000 times. Each permutation was generated by randomly assigning the 400 scans into groups of four and determining the corresponding subject specificity score.

The subjects that had all four scans in either the ‘improvement’ or ‘no improvement’ group were further examined to determine the regional variability, or lack thereof, in the RRF curves. The RRFs of model *B_r,s_* were averaged across all the subjects within the ‘improvement’ and ‘no improvement’ group separately. A second k-means clustering was performed on the estimated RRF curves at the voxel and parcel level to categorize the voxel and parcel-wise RRFs into groups with distinct dynamics. At the voxel level, only the gray matter was considered, and three clusters were used. Two clusters were used at the parcel level as this analysis did not include subcortical regions.

### 2.6. Association of PRF curves with vessel density, respiratory patterns and behavioral measurements

The relationship of the PRF curve features with arterial and venous densities obtained using the probabilistic maps of Bernier et al. (2018) was examined at the voxel and parcel level. At the parcel level, the vessel densities were averaged within each parcel. Furthermore, a weighted average was used to compute the mean PRF curve features across all subjects. The correlation between the two-input model and the parcel timeseries was used as a weighting coefficient to assign more weight to parcels of subjects that are more strongly influenced by physiological processes and, thus, were more reliable for extracting PRF curve features.

The parcel-wise differences between models *B_r_* and *B_p_*, calculated above, from all four scans for the 100 subjects were correlated with several respiratory measures from each scan. These included the standard deviation of the BR and RV signals, and the mean BR. Moreover, the differences between models *B_s_* and *B_p_* were averaged across the four scans of each subject and correlated across individuals with the age, weight, height, systolic blood pressure, diastolic blood pressure, body mass index (BMI) and hematocrit of each subject.

The arterial and venous densities were correlated with the following PRF curve features: FWHM, time-to-peak for the first and second peaks, as well as peak to height ratio. For the peak to height ratio, if the magnitude of the first peak was greater than the second peak, its value was assigned a positive value and if the magnitude of the second peak was greater it was assigned a negative value. In the cases where only one peak was observed, its value was set to 100 and made either positive or negative depending on the sign of the peak.

## 3. Results

### 3.1 Model evaluation

Figure 2 shows the performance of the examined models with respect to the BOLD variance explained at the voxel, parcel and whole-brain level. With regards to the one-input models, for the CRF we observe different results across spatial scales. At the voxel, parcel and whole brain levels, models *A_r_, B_p_* and *B_s_* yielded the best performance respectively. The RRF model exhibited a more consistent behavior across spatial scales, whereby the more flexible models using two gamma functions (*B_s_* and *B_r,s_*) yielded the best performance. Specifically, at the voxel and parcel levels, the best performance was yielded by model *B_r,s_*, while model *B_s_* performed best at the whole brain level. The same trend was observed in the behavior of the two-input model, i.e., model *B_r,s_* performed best at the voxel and parcel level and model *B_s_* performed best at the whole brain level. Overall, models using the basis set *B* performed better than models using the basis set *A,* despite the fewer degrees of freedom (2 vs 5 free parameters per input – HR and RV).

**Figure 2.**
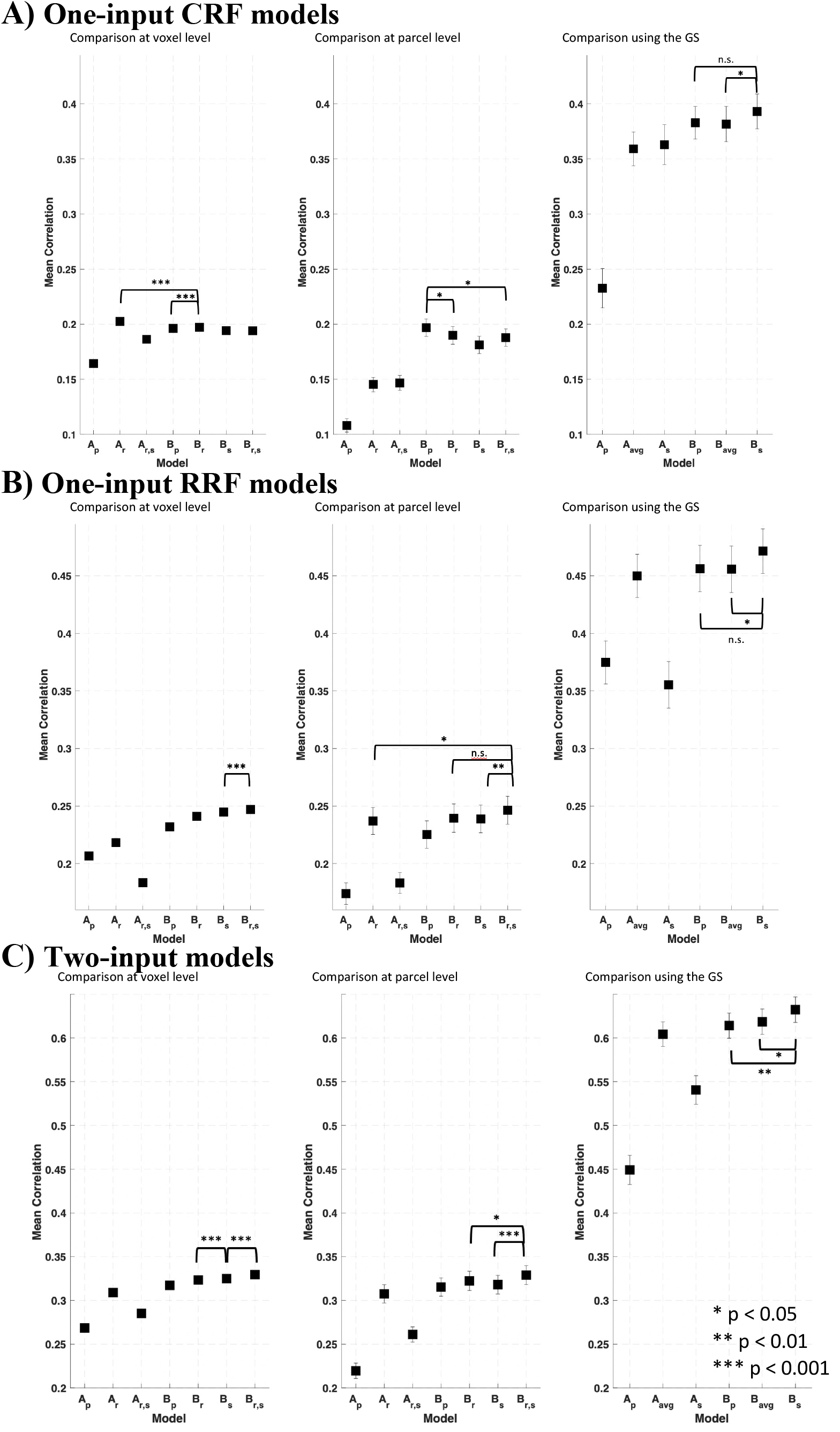
Comparison of model performance at different spatial scales (voxel, parcel and whole-brain level) for the cardiac component (A), respiratory component (B) and two-input model (C). Models *A_r_, B_p_* and *B_s_* yielded the best performance for the CRF at the voxel, parcel and whole brain level, respectively. Regarding the RRF, the best performan1ce1was yielded by the most flexible model using the basis set *B* (*B_r,s_* at the voxel and parcel level and model *B_s_* at the whole brain level). The same trend was observed for the two-input model as seen in the RRF where the best performance was yielded by the most flexible model using basis set *B* (*B_r,s_* at the voxel and parcel level and model *B_s_* at the whole brain level).

The seven models examined here yielded similar spatial maps of variance explained, with the highest correlation values observed in the gray matter. In addition, the top 5% of the voxels of each model corresponded mostly to voxels in the gray matter, as seen in Figure 3C, which shows a thresholded map averaged across all 100 subjects for model *B_r,s_*. Moreover, as evident from the correlation maps of the cardiac and respiratory models averaged across all 100 subjects (Figure 3A-B), variations in HR and RV affected similar regions, mainly in the visual and somatosensory cortex. The effects of respiration were generally found to be more pronounced as compared to HR.

**Figure 3.**
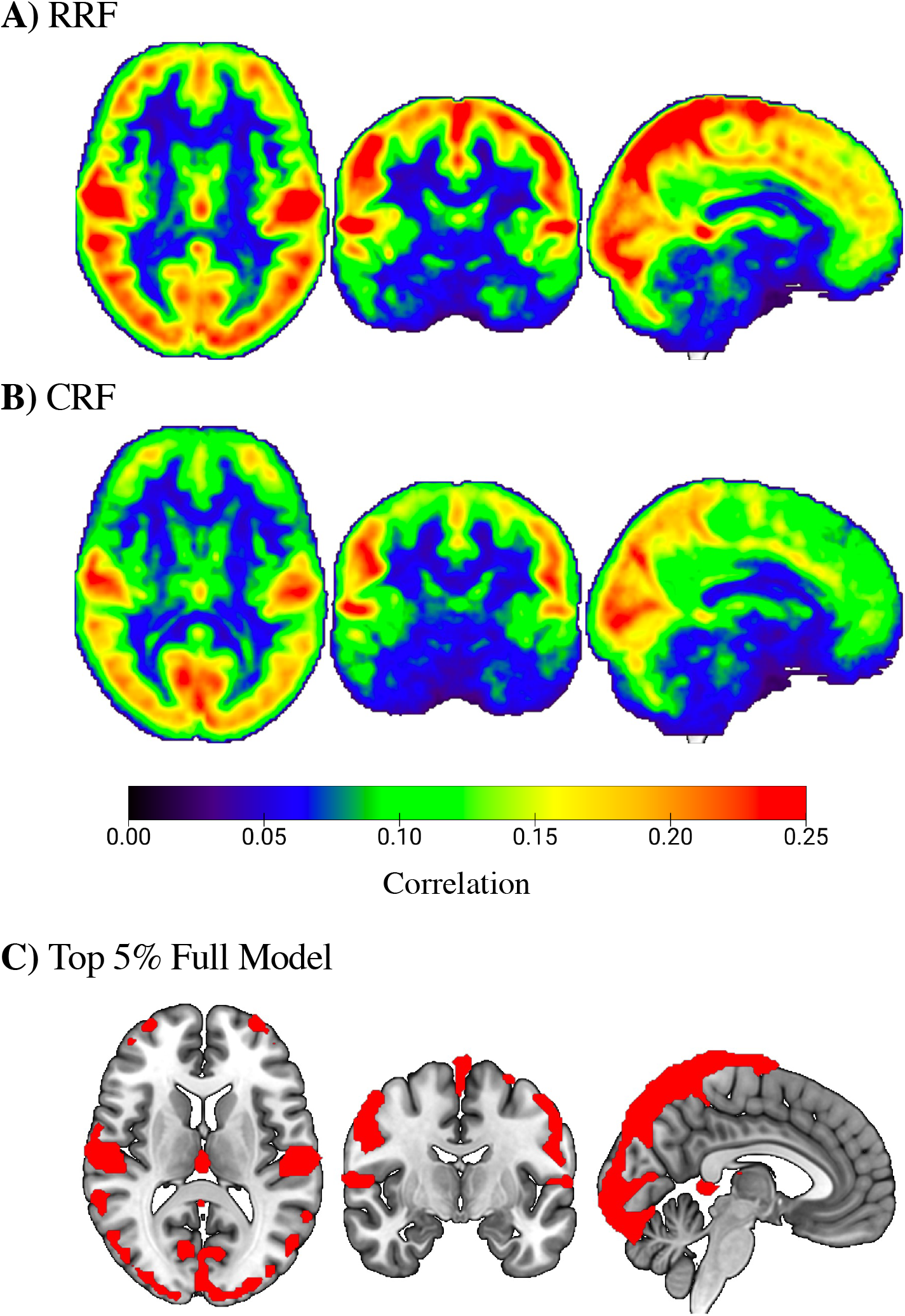
BOLD variance explained using model *B_r,s_*, averaged across all subjects (*N* = 100). A) Statistical map of the correlation between fMRI voxel time-series and RRF prediction. B) Statistical map of the correlation between fMRI voxel timeseries and CRF prediction. C) Mask with the voxels corresponding to the top 5% of correlation values obtained with the two-input model.

### 3.2. Assessment of regional and subject variability

Further analysis was carried out on the one-input RRF models using the basis set *B*. Specifically, the difference between the parcel-wise correlations of the RRF models *B_s_* and *B_p_* were further analyzed to investigate the advantage of allowing regional RRF variability. K-means clustering was performed to categorize the 400 scans from 100 subjects into two groups; a group that exhibited improvement when regional variability was allowed (‘improvement’ group) and a group that did not (‘no improvement’ group). Figure 4A shows the distribution of the differences in correlation values for each group across all 4 scans of the 100 subjects (4 scans x 100 subjects = 400 correlation values). The mean difference for the ‘improvement’ and ‘no improvement’ group were 0.042 and -0.006 respectively. Among the 400 scans, 37% fell into the ‘improvement’ group and 63% fell into ‘no improvement’ group. Figure 4B shows how the four scans of the 100 subjects were divided into the two groups. Among the 100 subjects, 38% had all four scans in the ‘no improvement’ group while 10% had all four scans in the ‘improvement’ group. Therefore, 48% of the subjects exhibited the same behavior across all four scans, which indicates a degree of subject specificity related to whether the use of a region-specific RRF improved performance. Of the remaining subjects, 15% had three out of four scans in the ‘no improvement’ group and 18% had three out of four scans in the ‘improvement’ group. This still indicates some degree of subject specificity related to whether the use of a region-specific RRF improved performance, as the majority of scans fell into the same group. Only 19% of subjects had the same number of scans in each group. To quantify this behavior, a subject specificity score was developed as described in Section 2.5, whereby subjects who had all scans in the same group were assigned a value of 1, subjects with one scan in one group and three scans in the other were assigned a value of 0.5, and subjects with two scans in one group and two in another were assigned a value of 0. The average subject specificity score across all subjects was found to be 0.65 with a standard deviation of 0.38. To validate the significance of this value, the score was calculated for 10,000 random permutations of the 400 scans. None of the permutation tests yielded a score equal or larger than 0.65, indicating strong evidence against the null hypothesis (*p* < 10^-4^).

**Figure 4.**
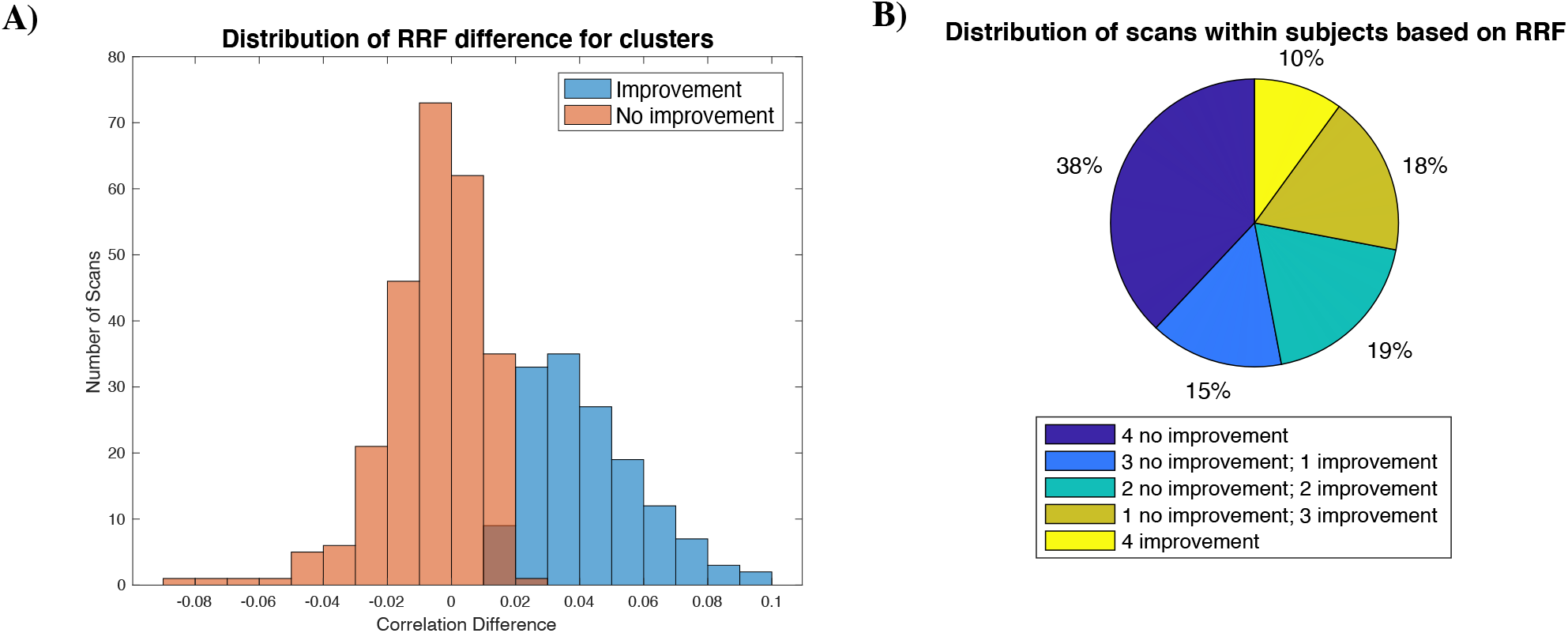
K-means clustering performed on the difference in correlation values obtained with the RRF models *B_p_* and *B*_*r*_. All four scans of each of the 100 subjects were considered. A) Histogram of the differences in correlation between the two models for all 400 scans, with the ‘improvement’ and ‘no improvement’ group indicated with orange and blue color respectively. B) Pie chart illustrating the distribution of scans across subjects according to the behavior with respect to using region-specific RRFs.

The RRFs of the two groups of subjects with either all scans in the ‘improvement’ group (10% of scans) or all scans in the ‘no improvement’ group (38% of scans) were further examined. The spatiotemporal RRF dynamics of these two groups averaged across all subjects within a group is shown at the parcel level in Figure 5 (the voxel-wise dynamics can be found in the Supp. Material, while videos of the dynamics can be found at https://doi.org/10.6084/m9.figshare.20715925.v2). The RRFs corresponding to these two groups exhibited different temporal dynamics for several brain regions. Specifically, the ‘improvement’ group yielded two spatially distinct clusters. Most regions yielded RRFs with negative peaks only, while the RRFs of the remaining regions exhibited a positive initial peak followed by a negative peak. In the ‘no improvement’ group, the dynamics were found to be more homogeneous spatially. Across all parcels, the RRFs exhibited a positive peak followed by a negative peak, with the amplitude of these peaks exhibiting some variability.

**Figure 5.**
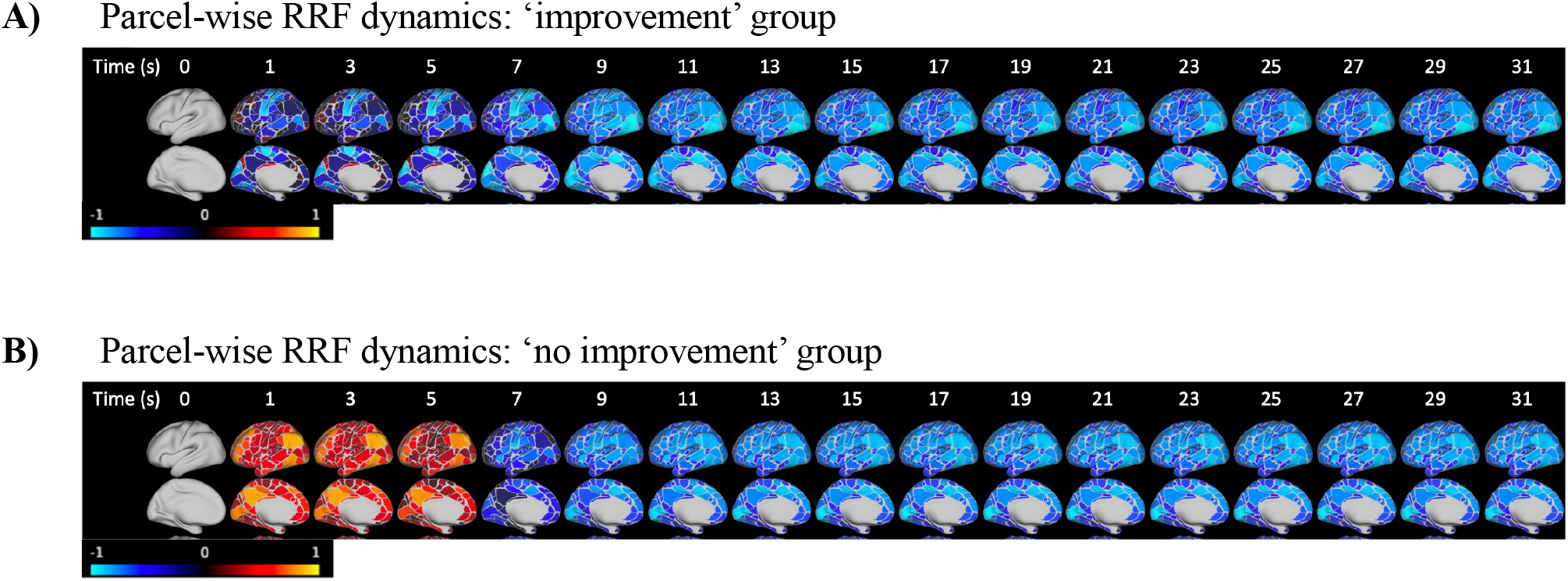
Spatiotemporal dynamics of the parcel-wise RRFs for the (A) ‘improvement’ group and the (B) ‘no improvement’ group. The former yielded a cluster of parcels with an initial positive peak and a cluster with an initial negative peak. Both clusters exhibited a secondary negative peak. The group with no improvement yielded a more spatially homogeneous response with a strong initial positive peak followed by a negative peak. Time corresponds to the time lag of the RRF in seconds.

The voxel and parcel-wise RRFs of *B_r,s_* from each group were averaged across all the subjects within each group and k-means clustering was performed on the curves to determine distinct groups of curve dynamics across brain regions. Two clusters were considered at the parcel level and three clusters at the voxel level, whereby the analysis in the latter case was restricted to voxels in the grey matter. Preliminary analysis evaluated the performance when using different numbers of clusters at the parcel level; however, using a higher number of clusters did not provide additional insight into the differences in regional RRF dynamics. An additional cluster was used at the voxel-level to account for the inclusion of sub-cortical regions. With regards to the voxel level analysis, for the ‘no improvement’ group the three clusters exhibited similar RRF curves with a positive peak followed by a stronger negative peak (Figure 6A). In contrast, for the ‘improvement’ group, the relative amplitude and polarity of the first peak exhibited distinct differences across the three clusters. Specifically, one cluster consisting mainly of cerebellum and brainstem regions (blue color) exhibited a large positive peak followed by a negative peak, while another cluster consisting of sensory-motor regions (red color) yielded two strong negative peaks, and a third cluster consisting of the remaining regions in the cortex (green color) exhibited a weak initial positive peak followed by a stronger negative peak. Similar trends were observed at the parcel level for the two identified clusters (Figure 6B). For instance, the cluster consisting of sensory-motor regions was characterized by two negative peaks for the ‘improvement’ group, while a positive peak followed by a negative peak was found in the ‘no improvement’ group.

**Figure 6.**
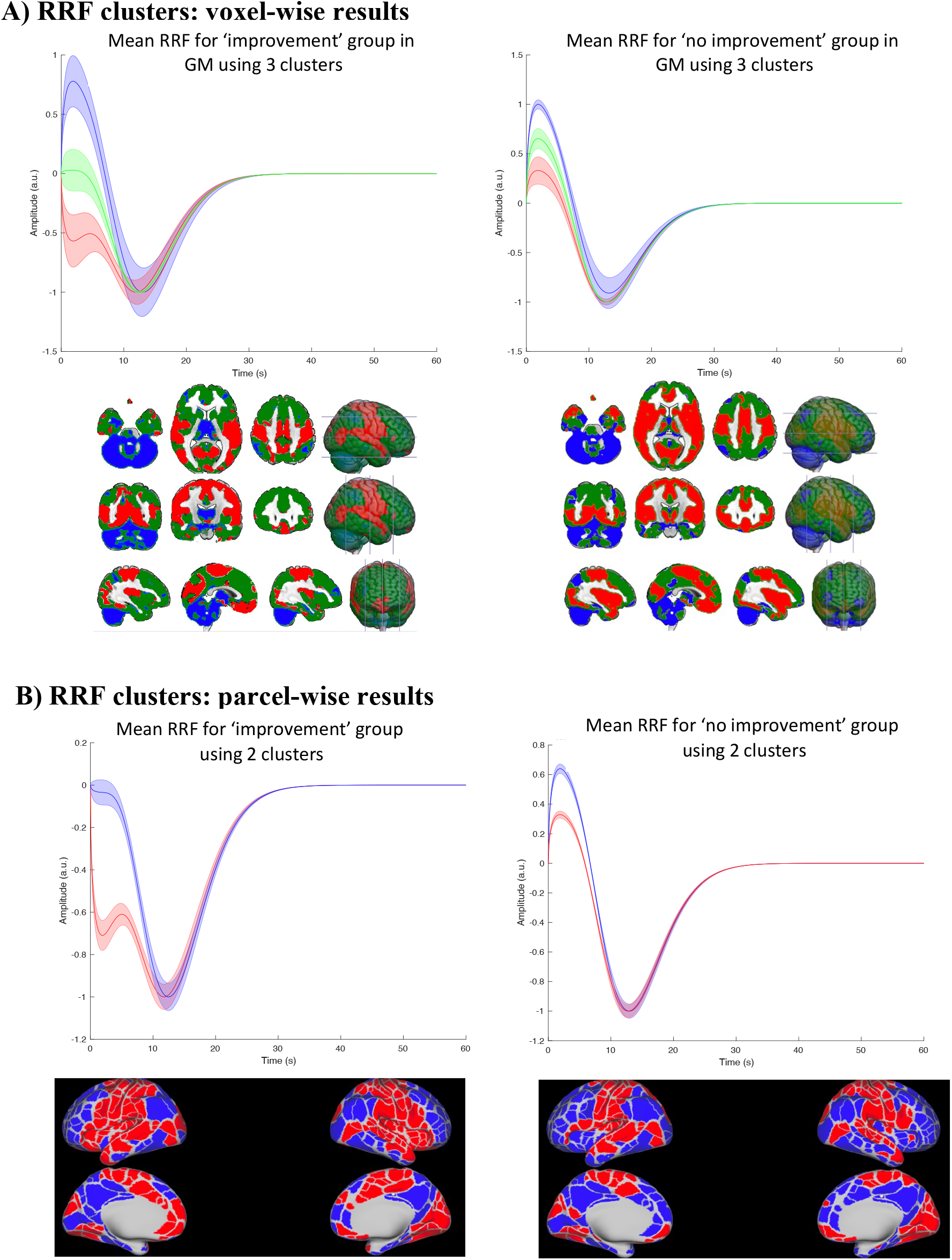
K-means clustering of RRF curve dynamics for the ‘improvement’ (left) and ‘no improvement’ (right) groups obtained at the (A) voxel and (B) parcel level (see Section 2.5). At the voxel level, three clusters were used and applied only to grey matter. The ‘no improvement’ group yielded clusters with variation only in their positive peak amplitude. The ‘improvement’ group yielded more variation in terms of RRF dynamics, with the blue cluster having a clear bimodal shape, the green cluster a very weak first peak followed by a negative peak and the red cluster two negative peaks. At the parcel level, the ‘no improvement’ group yielded two clusters with similar dynamics, albeit with small differences in the value of their positive peaks. The ‘improvement’ group yielded two distinct clusters of curve dynamics. Regions shown in blue color exhibited a weak initial peak followed by a negative peak, while regions shown in red color exhibited two negative peaks.

The spatial maps of the RRF of the parcels and the spatial maps of the voxels placed in each cluster for the ‘no improvement’ and ‘improvement’ groups exhibited similar patterns (Figure 6). The regions with an increased amplitude of the first peak were similar between the ‘improvement’ and ‘no improvement’ group. This trend was observed at both the parcel and voxel levels.

### 3.3 Association of PRF curves with vessel density, respiratory patterns and behavioral measurements

As described above, RRF features (e.g. FWHM of first peak) were extracted from the regionally specific PRFs obtained from the one-input models *B_s_* and correlated with the corresponding regional vessel density. At the parcel level, a weighted average was used to assign higher weight to parcels that exhibited better model performance, based on their correlation between the BOLD signal and the model prediction. The significance of each correlation was corrected for multiple comparisons (6 PRF curve features x 2 PRF curves x 2 vessel density maps = 24 tests). Once corrected for multiple comparisons, no correlations were found to be significant (Suppl. Table 1).

Finally, to determine why certain subjects or scans benefitted from allowing regional variability in RRF curves (*B_s_* vs *B*_*p*_), a series of tests was done to examine potential association of improvement with physiological features (i.e. body weight) as well as certain respiratory metrics (i.e. standard deviation of BR; see Section 2.6). However, no significant correlations were found (Suppl. Table 2, 3).

## 4. Discussion

We rigorously evaluated the performance of seven physiological models with varying complexity in modeling the dynamic effects of physiological processes (HR and RV) on BOLD fMRI signal fluctuations. All examined models utilized physiological response functions which were convolved with HR and RV. Some models assumed subject and spatial invariance for the PRF curves, eliminating the need for parameter fitting. On the other hand, for some models, PRF curves for each subject and/or region were estimated using least squares linear regression, allowing us to assess the extent to which using subject- and region-specific PRFs results in better prediction performance.

The mean PRF curves of the best performing model found in this study (*B_r,s_*; Figure 6) were, to a large degree, consistent with the dynamics reported in recent studies examining fMRI data from the HCP (Chen et al., 2020; Kassinopoulos & Mitsis, 2019, 2021). Interestingly, as previously noted, these PRF curves exhibited faster dynamics than the PRF curves reported in the early studies by Birn et al. (2008) and Chang et al. (2009).

As shown in Figure 2, when considering only the one-input CRF models, our results revealed no significant variation of CRF curve between brain regions and subjects. Conversely, allowing subject and regional variability for the RRF resulted in improved performance. Furthermore, results obtained using the two-input model suggested that allowing subject and regional variability for both the CRF and RRF leads to enhanced performance, but this improvement may stem from the significant improvement gained by allowing variability in the RRF, while being minimally affected by the variability in the CRF. Finally, it was found that the regional variability in the RRF yielded better performance for a subset of subjects (Figure 4).

### 4.1. Model evaluation

Our findings suggest that the RRF dynamics were more variable, regionally- and subject-wise, compared to the CRF dynamics, and this can be partly attributed to the presence of respiration-related motion artifacts. There is significant variability in respiration-related motion artefacts across subjects, as these artifacts depend on several factors such as the air volume changes in the lungs, the subject’s body type, their respiratory behavior and their position in the scanner (Byrge & Kennedy, 2018; Power et al., 2019; Raj et al., 2001). Further, respiration-related motion artefacts vary within a subject across voxels along the phase-encoding direction (Raj et al., 2001).

Despite having the highest number of degrees of freedom, model *A_r,s_* performed poorly compared to other models. This model uses five basis functions for the CRF and five basis functions for the RRF to allow regional and subject variability. It is possible that this flexibility caused overfitting of the model and the PRF curves estimated on the training dataset did not generalize well when applied to the testing dataset. Our results indicate that models using the basis set *B* that consists of two basis functions per input (HR and RV), may provide a better balance between flexibility and constraining the shapes of the curves, preventing overfitting.

### 4.2 Assessment of regional variability

The extended analysis of model *B_R,s_* provided further insight into the benefits of allowing regional variability in the RRF. The 400 examined scans were clustered into a group of scans which exhibited improvement when regional variability was allowed in the one-input RRF model, and a group which did not exhibit improvement (Figure 4). Nearly 50% of the subjects had all four of their scans clustered into the same group (i.e. ‘improvement’ and ‘no improvement’), while a further 30% had three out of four scans in the same cluster. This behavior was also reflected in the subject specificity score, which was found to be significantly higher than chance level. These results suggest that allowing regional variability in RRF is beneficial only for specific subjects; however, we were not able to identify any specific factors that could explain this behavior.

To examine the different behavior in RRF dynamics between the ‘improvement’ and ‘no improvement’ groups, the RRFs of model *B_r,s_* were averaged within the two groups. The RRF curves obtained from all voxels were clustered within each group to investigate the differences in PRF dynamics across the brain. The subject group with improvement exhibited three distinct clusters of PRF curve dynamics (Figure 6); some regions exhibited the typical RRF curve with a positive peak preceding a negative peak (e.g. visual cortex), while other regions only exhibited the second negative peak (e.g. frontal cortex) or two distinct negative peaks (e.g. somatosensory regions). Conversely, the ‘no improvement’ group yielded variation in the amplitude of the first peak only (videos of the aforementioned voxel-wise dynamics can be found at https://doi.org/10.6084/m9.figshare.20715925.v2).

The benefits of allowing regional variability for the ‘improvement’ group are mainly due to that the RRF curves in various regions, including the somatosensory regions, exhibited an atypical RRF curve with two distinct negative peaks. A potential explanation for this atypical RRF curve could be that these subjects were more aware of their respiratory patterns, perhaps due to anxiety caused by lying inside the MR scanner, leading to a condition known as respiratory interoception. Respiratory interoception may involve regions beyond the ones implicated in previous studies (Harrison et al. 2021), and the underlying hemodynamic response associated with this elevated interoception may somehow interfere with the detection of the RRF curve. Further investigations are needed to examine the role of respiratory interoception in the estimation of RRF curves.

### 4.3. Association of RRF curves with physiological factors

To better understand why region-specific RRF yielded improved performance only for some subjects, we examined whether the extent of improvement achieved when allowing regional variability was associated to inter-individual differences in physiological features (e.g. body weight) or respiratory metrics (e.g. variations in BR). However, none of the tests yielded statistically significant results. In addition, we examined the relationship between vascular density and RRF shape. More precisely, we tested whether regional differences in PRF shape across parcels correlated with regional differences in vascular density. However, once again, none of the correlations was significant.

The low correlations found at the parcel level were not entirely unexpected (Suppl. Table 1), considering the significant variability between subjects with respect to their vascular density profiles (Bernier et al., 2018). It has been suggested that the inter-subject variability in vascular density profiles increases with decreasing vessel size (Bernier et al., 2018). Moreover, as the effects of heart rate and respiratory variations are more pronounced in venules rather than large draining veins, it is possible that subject-specific vessel density maps are necessary to detect any relationships between RRF curves with vascular density profiles.

Chen et al. (2020) found a relationship between the variability of vascular density in different brain regions and RRF/CRF amplitudes, although their analysis was strictly qualitative. Furthermore, it is plausible that the shape of the PRF curves is also likely influenced by the path that blood takes throughout the brain. Future research incorporating improved characterization of vascular anatomy at the subject level is needed to better understand the dependence of regional PRF curves on vascular density.

### 4.4. Limitations

This study only investigated resting state scans only, and the results may not be directly applicable to task-based fMRI studies. The tasks implemented in these studies typically require participants to maintain relatively stable levels of vigilance. Consequently, arousal levels may not modulate cardiac and respiratory activity as strongly as observed in resting-state fMRI. However, some tasks may change the mean value and variability of physiological signals (heart rate, respirations), which would elicit changes in the BOLD fMRI timeseries, and overlap with the neural-related BOLD activity associated to the task. In this context, conducting simultaneous EEG-fMRI studies could provide additional insights into the interactions between physiologically- and neurally-driven BOLD responses during task-based paradigms.

Furthermore, an additional limitation of the current and previous relevant studies is the assumption of stationarity in PRF curves over time. The performance of the models that required estimation of PRF curves on a subject-specific basis, was evaluated by training them using the first of two scans collected in a session and testing them using the second scan of the session. This cross-validation scheme assumes that the PRF curves are stable within a subject within the span of an hour. However, we acknowledge that the PRF curves may vary across scans of the same session (i.e. scans collected briefly one after the other), and future research should examine this possibility.

## 5. Conclusions

We demonstrated that incorporating regional and subject variability in the PRF curves improves the ability of linear PRF one-input and two-input dynamic models to explain BOLD variance induced by physiological fluctuations. This improvement was particularly noticeable for respiration, and especially for a subset of subjects who exhibited a wider range of PRF dynamics compared to other subjects. These insights pave the way for more effective removal of cardiac and respiratory fluctuations from fMRI recordings, thereby facilitating the disentanglement of the underlying neural activity from physiological confounds.

## Supplementary Material

### Subject List

|  |  |  |  |  |  |  |
| --- | --- | --- | --- | --- | --- | --- |
| '100307' | '100610' | '105115' | '107321' | '108121' | '113922' | '117930' |
| '118528' | '118730' | '118932' | '119732' | '120212' | '120717' | '121416' |
| '121618' | '121921' | '122620' | '122822' | '123521' | '123824' | '128127' |
| '128632' | '132118' | '137027' | '138837' | '139435' | '154532' | '154734' |
| '158843' | '159239' | '159441' | '159744' | '160830' | '172938' | '191437' |
| '192136' | '196346' | '200008' | '200614' | '200917' | '201414' | '201818' |
| '203418' | '203923' | '204622' | '207123' | '207426' | '208630' | '209127' |
| '210011' | '210415' | '210617' | '300618' | '300719' | '304020' | '305830' |
| '307127' | '309636' | '393247' | '395251' | '395756' | '395958' | '406432' |
| '412528' | '421226' | '422632' | '424939' | '441939' | '520228' | '536647' |
| '555348' | '580751' | '581450' | '585256' | '588565' | '592455' | '599065' |
| '654350' | '656657' | '668361' | '685058' | '686969' | '690152' | '693764' |
| '694362' | '700634' | '704238' | '800941' | '833249' | '852455' | '878776' |
| '880157' | '894774' | '896778' | '901442' | '910241' | '911849' | '917558' |
| '923755' | '926862' |  |  |  |  |  |

## Results

**Suppl. Figure 1.**
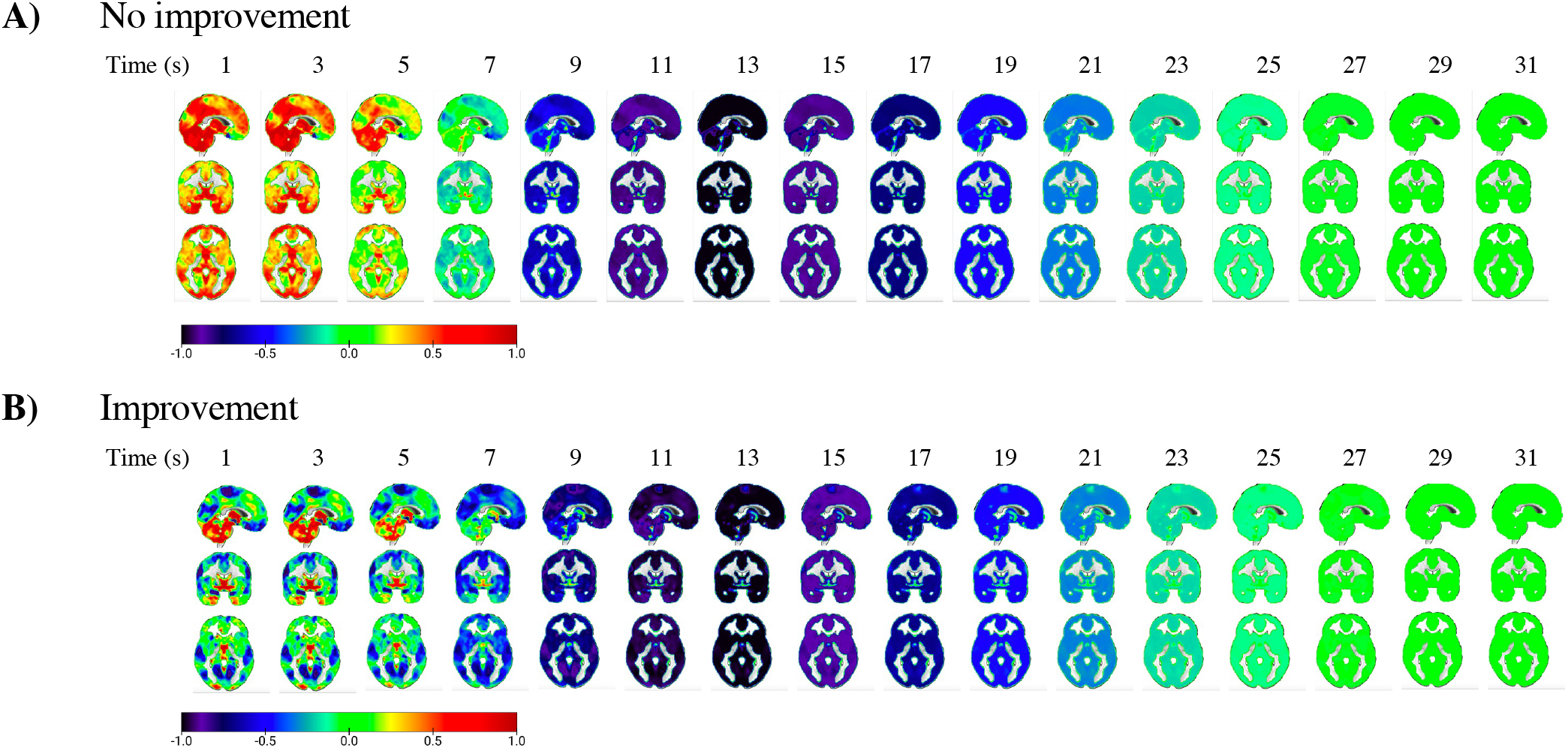
Spatiotemporal dynamics of the voxel-wise RRFs for the (A) group with improvement and the (B) group with no improvement. The group with improvement has a cluster of voxels with an initial positive peak and another cluster of voxels with an initial negative peak. Both clusters then have a secondary negative peak. The group with no improvement has a more spatially homogeneous response with a strong initial positive peak followed by a negative peak.

**Suppl. Table 1.**
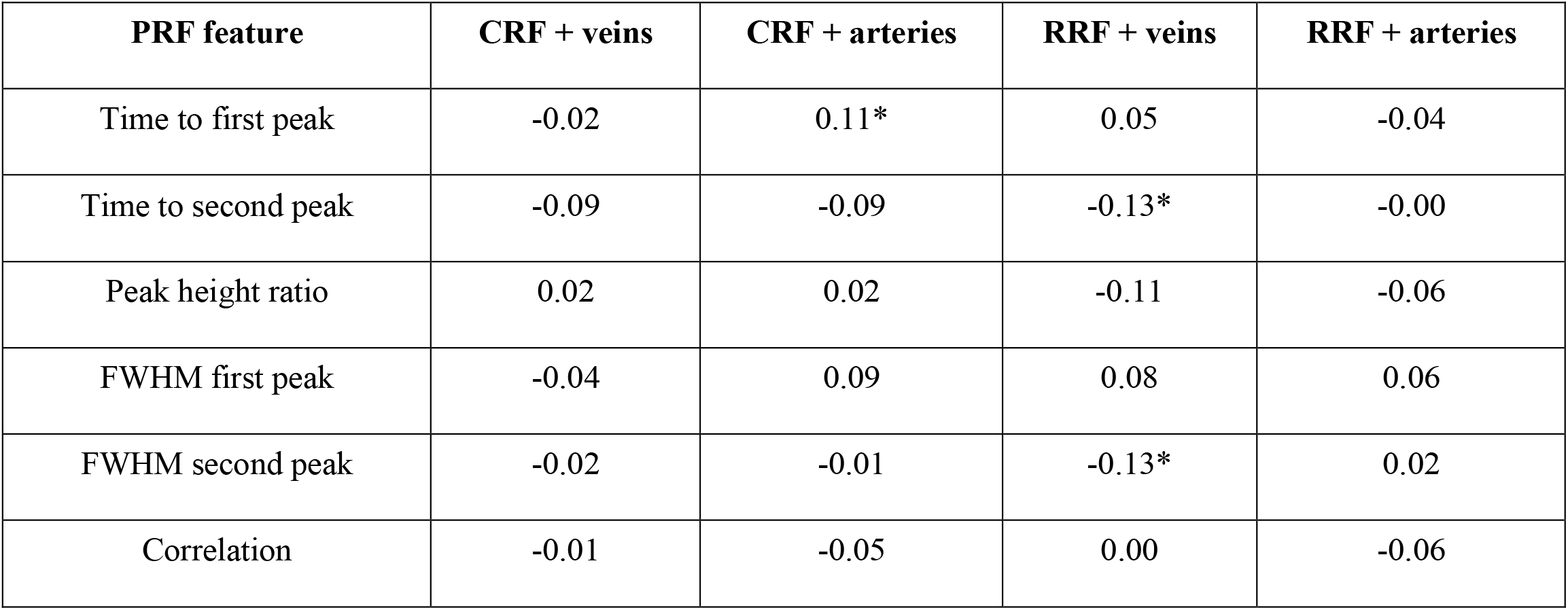
Correlations of PRF features with vessel densities at the parcel level. * p < 0.05.

| PRF feature | CRF + veins | CRF + arteries | RRF + veins | RRF + arteries |
| --- | --- | --- | --- | --- |
| Time to first peak | -0.02 | 0.11* | 0.05 | -0.04 |
| Time to second peak | -0.09 | -0.09 | -0.13* | -0.00 |
| Peak height ratio | 0.02 | 0.02 | -0.11 | -0.06 |
| FWHM first peak | -0.04 | 0.09 | 0.08 | 0.06 |
| FWHM second peak | -0.02 | -0.01 | -0.13* | 0.02 |
| Correlation | -0.01 | -0.05 | 0.00 | -0.06 |

**Suppl. Table 2.** Correlations between respiratory metrics and voxel-wise model performance difference between *B_r_* and *B_p_*. None of the correlations was significant (p<0.05).

| **Respiratory Metric** | **Correlation** |
| --- | --- |
| RV standard deviation | 0.07 |
| BR standard deviation | -0.06 |
| Mean BR | 0.02 |

**Suppl. Table 3.** Correlations between physiological metrics and voxel-wise model performance difference between *B_r_* and *B_p_*. None of the correlations was significant (p<0.05).

| **Measurement** | **Correlation** |
| --- | --- |
| Age | 0.14 |
| Height | 0.11 |
| Weight | 0.17 |
| BMI | -0.15 |
| BP systolic | -0.07 |
| BP diastolic | -0.11 |
| Hematocrit | -0.11 |

